# Mutational spectra of SARS-CoV-2 orf1ab polyprotein and Signature mutations in the United States of America

**DOI:** 10.1101/2020.05.01.071654

**Authors:** Shuvam Banerjee, Sohan Seal, Riju Dey, Kousik Kr. Mondal, Pritha Bhattacharjee

## Abstract

Pandemic COVID-19 outbreak has been caused due to SARS-COV2 pathogen, resulting millions of infection and death worldwide, USA being on top at the present moment. The long, complex orf1ab polyproteins of SARS-COV2 play an important role in viral RNA synthesis. To assess the impact of mutations in this important domain, we analyzed 1134 complete protein sequences of orf1ab polyprotein from NCBI Virus database from affected patients across various states of USA from December 2019 to 25^th^ April, 2020. Multiple sequence alignment using Clustal Omega followed by statistical significance was calculated. Four significant mutations T265I (nsp 2), P4715L (nsp 12) and P5828L and Y5865C (both at nsp 13) were identified in important non-structural proteins, which function either as replicase or helicase. A comparative analysis shows 265T>I, 5828P>L and 5865Y>C are unique to USA and not reported from Europe or Asia; while one, 4715P>L is predominant in both Europe and USA. Mutational changes in amino acids are predicted to alter structure and function of corresponding proteins, thereby it is imperative to consider the mutational spectra while designing new antiviral therapeutics targeting viral orf1ab.

## Introduction

SARS-CoV-2 is the responsible pathogen for pandemic COVID 19. Positive-stranded, RNA genomes of Coronaviruses is around 27 to 32-kb in length, of which about 2/3^rd^ encompasses viral Orf1ab gene and expresses the largest and most complex polyproteins of any RNA viruses. The open reading frame 1 (ORF1), functions as replicase, replicase/transcriptase, or polymerase polyproteins, is translated into ORF1a (approximately 486kDa, major product) and ORF1b (∼306KDa) polyproteins in the host cell. Virus-encoded proteinases including Papain like protease (PLPs) and 3C like protease (3CL Pro) cleaves ORF1 into 16 nonstructural proteins (nsps)(1,2). ORF1a comprising nsps (nsp 1 to nsp 10) play an important role in coping cellular stresses and maintaining functional integrity of the cellular components along with the pivotal roles in viral replication. On the other hand, ORF1b encodes viral RNA-dependent RNA polymerase (nsp12), helicase (nsp13), exonuclease (nsp14), a polyU (Uridylate) specific endonuclease (nsp15), and methyl transferase (nsp16). Hence, majority of these nsps play an important role in viral pathogenesis and promising target for anti-viral drug targeting and vaccine synthesis (3).

At present, USA is one of the worst affected countries globally in terms of affected individuals and the number of death, is concerned. Till 25^th^ April, USA is reported to have 860,772 positive cases and 44,053 deaths (4). This adverse condition led to investigate the sequence of viral whole genome reported to NCBI virus database. As on 25^th^ April 2020 (till 12 noon, IST), around 1134 Orf1ab polyprotein sequence have been submitted from USA alone. Different states of USA like Washington D.C, New York, Connecticut, Idaho, Georgia etc have uploaded sequence of Orf1ab polyprotein in the database. Since COVID-19 originated in Wuhan and found to be extended to different parts of the globe with variation in its virulence, it is imperative to identify the mutations occurred in Orf1ab polyprotein and consequent impact in protein structure and interaction with the host body. Hence the present study aims (i) to identify the mutations observed in orf1ab polyprotein, (ii) to predict the conformational changes of SARS-CoV-2 polyprotein due to the mutations and (iii) to identify the signature pattern, if any for USA.

## Methodology

### ORF1ab protein sequence retrieval from the database

The protein sequences were retrieved from “NCBI Virus” database, specific input was “SARS-CoV-2”. Then the output was refined with sequence length 7050 to 7100, as the length of our target orf1ab polyprotein is 7096. Total 1307 sequences were retrieved, among them 125 sequences were from Asia, 42 from Europe and 1134 solely from USA.

### Screening of submitted sequences and selection of study sample

Incomplete sequences or sequence with undetermined residues (mentioned as X) were eliminated. A total of 867 orf1ab polyprotein complete sequences deposited from 31 different states of USA were considered for the present study (Group A). All the above-mentioned sequences from Asia (Group B) and Europe (Group C) were also selected.

### Multiple Sequence Alignment (MSA) and analysis of mutational spectra

Clustal Omega (5) was employed to align multiple sequences of each of the above-mentioned groups. Then sequences of Group A were subdivided into 31 subgroups depending on the regional source from which the sequence was originated and MSA was conducted encompassing all subgroups using ancestral orf1ab polyprotein sequence of Wuhan (YP_009742608, comprising 7096 residues) as reference sequence (6,7). Gap opening penalty and gap extension penalty was set 12 and 2 respectively to ensure that unnecessary gaps are not created during alignment and alterations are visualised easily. Alignment results were thoroughly screened to find out the exact location of the mutations and alteration linked to that position.

### Calculation of statistical significance to detect signature mutations of USA

The number of occurrence of each mutated variant in Group A, B and C were calculated (suppose a, b and c) and then divided by the total number of sequence submitted under that group (suppose x for Group A, y for Group B and z for Group C). The proportion of each variant in Group A would be a/x and likewise for other Groups. Then p-value was calculated through Z-score using the proportion of the mutant and sample size to establish whether the occurrence of that mutant variant in USA is significantly higher in comparison to Asia and Europe. Two-tailed p values were calculated using 0.05 as the significance level (8). Thus attempt has been made to identify the signature mutations in the region of orf1ab polyprotein.

### Homology Modelling and simulation of protein structure

Structures of the associated non structural proteins for wild type were reported at I-Tasser server, however, the structures were not available for the varied amino acid alteration. To identify the alteration, Homology Modelling method by I-Tasser was used to generate secondary structure, which was then superimposed with the wild type using UCSF Chimera and PyMOL for easy visualization and comparison.

## Results and Discussion

To assess the structural variation and identify the signature mutations, if any among the viral strain(s) identified from USA, all 1134 sequences, submitted to NCBI Virus database (December 2019 to 25^th^ April 2020) were retrieved. Following the exclusion criteria mentioned above, 867 complete sequences from 31 different states of USA were obtained. For each subgroup of Group A, mutations observed in >5% studied population were taken into considerations and shown in **Table 1**.

**Table 1.** Region wise list of mutations at orf1ab polyprotein in USA.

| Name of the State | Sample size | Mutation Position | Location in orf1ab | Amino acid alteration | Individual with mutated variant | % carrying mutated variant |
| --- | --- | --- | --- | --- | --- | --- |
| Washington | 500 | 265 | nsp 2 | T > I | 206 | 41.2 |
|  |  | 4715 | nsp12 | P > L | 226 | 45.2 |
|  |  | 5828 | nsp 13 | P > L | 255 | 51.0 |
|  |  | 5865 | nsp 13 | Y > C | 255 | 51.0 |
| New York | 150 | 265 | nsp 2 | T > I | 114 | 76.0 |
|  |  | 3884 | nsp 7 | S > L | 28 | 18.7 |
|  |  | 4715 | nsp 12 | P > L | 135 | 90.0 |
|  |  | 6245 | nsp 14 | A > V | 24 | 16 |
| Virginia | 52 | 265 | nsp 2 | T > I | 17 | 32.7 |
|  |  | 3483 | nsp 5 | L > F | 5 | 9.6 |
|  |  | 4715 | nsp 12 | P > L | 38 | 73.1 |
|  |  | 5828 | nsp 13 | P > L | 10 | 19.2 |
|  |  | 5865 | nsp 13 | Y > C | 10 | 19.2 |
| Idaho | 21 | 265 | nsp 2 | T > I | 18 | 85.7 |
|  |  | 4715 | nsp 12 | P > L | 19 | 90.5 |
| Connecticut | 17 | 265 | nsp 2 | T > I | 11 | 64.7 |
|  |  | 4715 | nsp 12 | P > L | 14 | 82.4 |
|  |  | 5828 | nsp 13 | P > L | 1 | 5.9 |
|  |  | 5865 | nsp 13 | Y > C | 1 | 5.9 |
| California | 16 | 265 | nsp 2 | T > I | 4 | 25.0 |
|  |  | 4715 | nsp 12 | P > L | 5 | 31.3 |
| Massachusetts | 16 | 4715 | nsp 12 | P > L | 15 | 93.8 |
| Georgia | 13 | 971 | nsp 3 | P > L | 8 | 61.5 |
|  |  | 3606 | nsp 6 | L > F | 2 | 15.3 |
|  |  | 4715 | nsp 12 | P > L | 3 | 23.1 |
|  |  | 6158 | nsp 14 | F > L | 8 | 61.5 |
| Utah | 10 | 265 | nsp 2 | T > I | 4 | 40 |
|  |  | 4715 | nsp 12 | P > L | 5 | 50 |
|  |  | 5828 | nsp 13 | P > L | 4 | 40 |
|  |  | 5865 | nsp 13 | Y > C | 4 | 40 |
| Minnesota | 8 | 265 | nsp 2 | T > I | 3 | 37.5 |
|  |  | 4715 | nsp 12 | P > L | 3 | 37.5 |
|  |  | 5828 | nsp 13 | P > L | 3 | 37.5 |
|  |  | 5865 | nsp 14 | Y > C | 3 | 37.5 |
| Florida | 8 | 3606 | nsp 6 | L > F | 1 | 12.5 |
|  |  | 4715 | nsp 12 | P > L | 5 | 62.5 |
|  |  | 5828 | nsp 13 | P > L | 1 | 12.5 |
|  |  | 5865 | nsp 13 | Y > C | 1 | 12.5 |
| Illionis | 7 | 265 | nsp 2 | T > I | 1 | 14.3 |
|  |  | 4715 | nsp 12 | P > L | 3 | 42.8 |
|  |  | 5828 | nsp 13 | P > L | 1 | 14.3 |
|  |  | 5865 | nsp 13 | Y > C | 1 | 14.3 |
| Iowa | 7 | 4715 | nsp 12 | P > L | 7 | 100 |
| Pennsylvania | 7 | 4715 | nsp 12 | P > L | 7 | 100 |
| New Hampshire | 4 | 265 | nsp 2 | T > I | 3 | 75 |
|  |  | 4715 | nsp 12 | P > L | 4 | 100 |
|  |  | 4764 | nsp 12 | L > F | 1 | 25 |
| N. Carolina | 4 | 265 | nsp 2 | T > I | 1 | 25 |
|  |  | 4715 | nsp 12 | P > L | 1 | 25 |
|  |  | 5828 | nsp 13 | P > L | 1 | 25 |
|  |  | 5865 | nsp 13 | Y > C | 1 | 25 |
| Texas | 3 | 2124 | nsp 3 | T > I | 2 | 66.7 |
|  |  | 3606 | nsp 6 | L > F | 2 | 66.7 |
|  |  | 4715 | nsp 12 | P > L | 2 | 66.7 |
| Arizona | 3 | 265 | nsp 2 | T > I | 1 | 33.3 |
|  |  | 4715 | nsp 12 | P > L | 1 | 33.3 |
|  |  | 5828 | nsp 13 | P > L | 2 | 66.7 |
|  |  | 5865 | nsp 13 | Y > C | 2 | 66.7 |
| Ohio | 3 | 3606 | nsp 6 | L > F | 1 | 33.3 |
|  |  | 4715 | nsp 12 | P > L | 2 | 66.7 |
| Rhode Island | 3 | 265 | nsp 2 | T > I | 1 | 33.3 |
|  |  | 4715 | nsp 12 | P > L | 3 | 100 |
| Nebraska | 2 | 392 | nsp 2 | G > D | 2 | 100 |
|  |  | 876 | nsp 3 | A > T | 2 | 100 |
|  |  | 4715 | nsp 12 | P > L | 2 | 100 |
| Nevada | 2 | 3606 | nsp 6 | L > F | 1 | 50 |
|  |  | 6669 | nsp 15 | A > V | 1 | 50 |
| S. Carolina | 2 | 971 | nsp 3 | P > L | 2 | 100 |
|  |  | 5401 | nsp 13 | P > L | 1 | 50 |
|  |  | 6158 | nsp 14 | F > L | 2 | 100 |
| New Orlando | 2 | 265 | nsp 2 | T > I | 2 | 100 |
|  |  | 4715 | nsp 12 | P > L | 2 | 100 |
| Kansas | 1 | 4715 | nsp 12 | P > L | 1 | 100 |
| Louisiana | 1 | 265 | nsp 2 | T > I | 1 | 100 |
|  |  | 3352 | nsp 5 | L > F | 1 | 100 |
|  |  | 4715 | nsp 12 | P > L | 1 | 100 |
| Maryland | 1 | 4715 | nsp 12 | P > L | 1 | 100 |
| Missouri | 1 | 4715 | nsp 12 | P > L | 1 | 100 |
| New Jersey | 1 | 4715 | nsp 12 | P > L | 1 | 100 |
|  |  | 6528 | nsp 15 | G > D | 1 | 100 |
| Hawaii | 1 | 4715 | nsp 12 | P > L | 1 | 100 |
| Wisconsin | 1 | 6679 | nsp 15 | L > del | 1 | 100 |
| Total | 867 |  |  |  |  |  |

Mutation at amino acid (AA) location 265 (Thr > Ile) of nsp2 is observed among ∼50% states (subgroups) and 44.2% sequences in USA (Table 1). Nsp2 is an important domain that takes care of the functional integrity of mitochondria and copes up cellular stress (9). In addition to this, nsp2 co-operates with nsp4 in viral replication. Nsp12 is important for its RNA dependent RNA polymerase (RdRp) activity and Pro > Leu in 4715 at this domain is a significant alteration, reported in 28 out of 31 states. Nsp 13 functions as helicase protein, which is a key enzyme in viral replication and therefore, two mutations, P5828L and Y5865C observed in this domain, expected to alter in structure-function and interaction with host’s target site. These two mutations, in particular are observed and presumed to be linked and found in equal proportions (0.321), among affected individuals from 9 out of 31 states.

All these four mutations are widely observed throughout the country but some mutations are consistently observed to specific regions (**Figure 1**). Georgia and South Carolina are two neighbouring states sharing common border in the South East region of USA and both carry mutant variant Leucine at the position of 971 (P>L) at nsp 3 and 6158 (F>L) at nsp14 in more than 60% cases. We found another mutation 3606L>F in nsp6 and individuals from Florida, Texas, Ohio, Nevada and Georgia had this mutation with a frequency of >10%. Nsp 6 plays a role in initial induction of autophagosomes from host endoplasmic reticulum, but gradually limits the expansion of these phagosomes which are unable to deliver viral components to lysosomes. It is important to note here that 3606L>F mutations were found in Italy while analyzing Group C (present study), which further strengthens our previous reporting (10).

**Figure 1.**
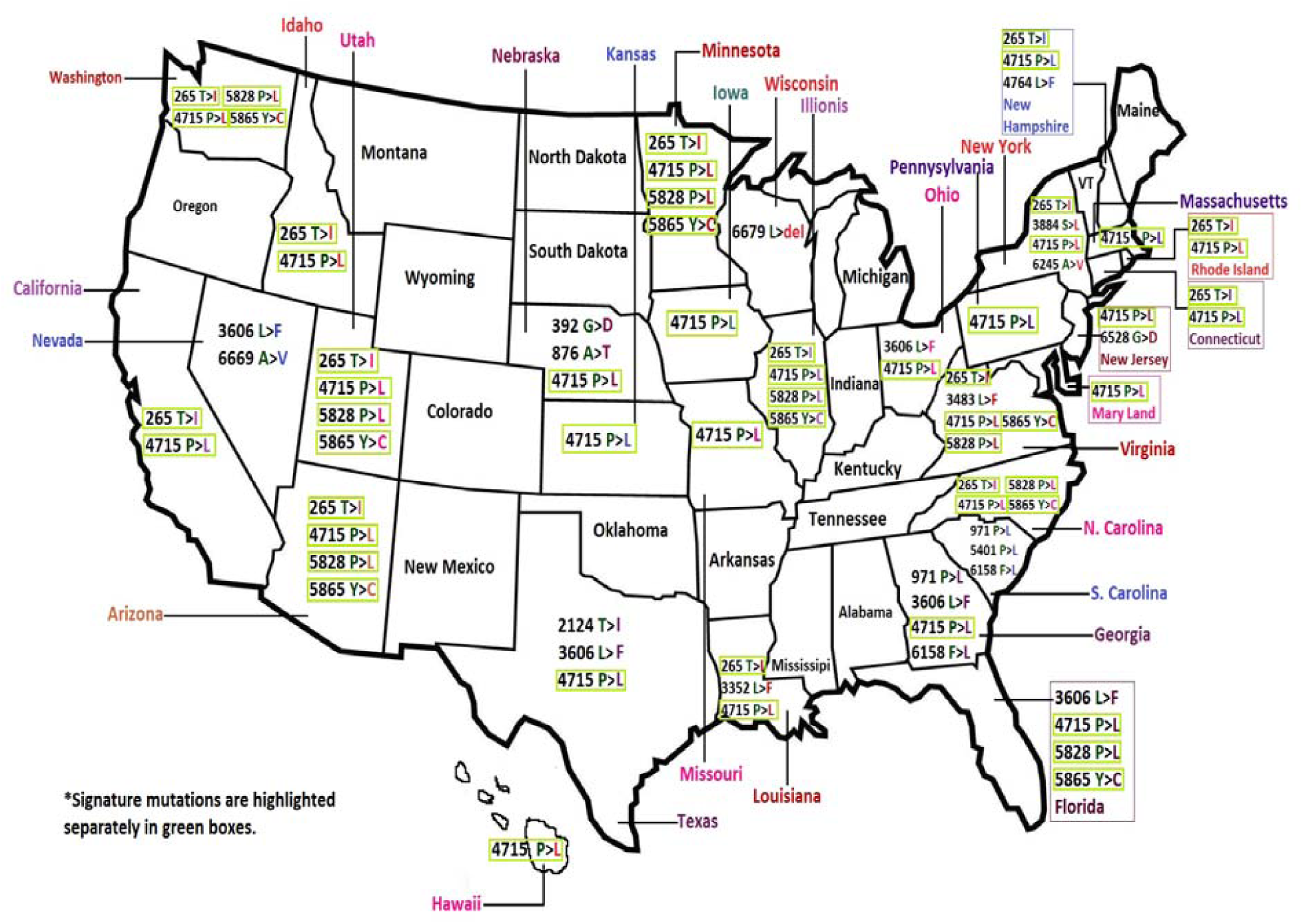
Mutational hotspots and the corresponding variants found in different geographical regions of USA

To study the implication of these mutations, we have compared the prevalence of T265I in European and Asian population with that of USA population. Strikingly, Threonine at 265 is the wild type AA found in all sequences from Europe and 119 out of 125 sequences from Asia. Therefore, the mutated AA Isoleucine is present in only 4.8% patient population in Asia, null in European population so far, but observed in 44.2% patient population (383 out of 867) in USA establishing it to be a signature mutation of the country (p = <0.0001).

Let us consider the mutation 4715 P > L, which is quite common in Europe and present in countries like Spain, France, Greece etc. and the amino acid variant Leucine is found in 51.6% (16 out of 31 sequences deposited by European countries). In USA, the frequency of Leucine is 58.1%, not significantly higher in comparison to Europe (p = 0.47) but clearly compared to Asia (<0.0001). However, the mutant Leucine at 4715 is consistently observed in America. Two other most frequent mutations in USA are 5828 P > L and 5865 Y > C, found in equal proportion (both observed 278 out of 867), presumed to be linked, and exclusively found in USA, so far (**Table 2**).

**Table 2.** The signature mutations of USA and proportional presence of those mutations in Asian and European continent

| Mutational hotspot in orflab pp | Location in orflab pp | USA (Grp 1) | Asia (Grp 2) | p-value (Grp 1 and Grp 2) | Europe (Grp 3) | p-value (Grp 1 and Grp 3) |
| --- | --- | --- | --- | --- | --- | --- |
| T265I | nsp 2 | 383/867<br>= 0.442 | 6/125<br>= 0.048 | <0.00001 | 0/31<br>= 0.0 | <0.00001 |
| P4715L | nsp 12 or RdRp | 504/867<br>= 0.581 | 14/125<br>= 0.112 | <0.00001 | 16/31<br>= 0.516 | 0.47152 |
| P5828L | nsp 13 or Helicase | 278/867<br>= 0.321 | 0/125<br>= 0 | <0.00001 | 0/31<br>= 0.0 | 0.00014 |
| Y5865C | nsp 13 or Helicase | 278/867<br>= 0.321 | 0/125<br>= 0 | <0.00001 | 0/31<br>= 0.0 | 0.00014 |

The four signature mutations are located in nsp 2, nsp 12 and nsp 13 respectively (**Figure 2**). Non-structural protein 2 is assumed to be responsible in the modulation of host cell survival signaling pathway by interacting with PHB and PHB2 in the host body (9, 11, 12). Change of a polar amino acid (Threonine) to a non-polar one (Isoleucine) makes it hydrophobic and induces structural alteration in that domain which is observed by simulating structure of the nsp2 protein harboring mutated allele through homology modeling using I-Tasser (13, 14, 15).

**Figure 2.**
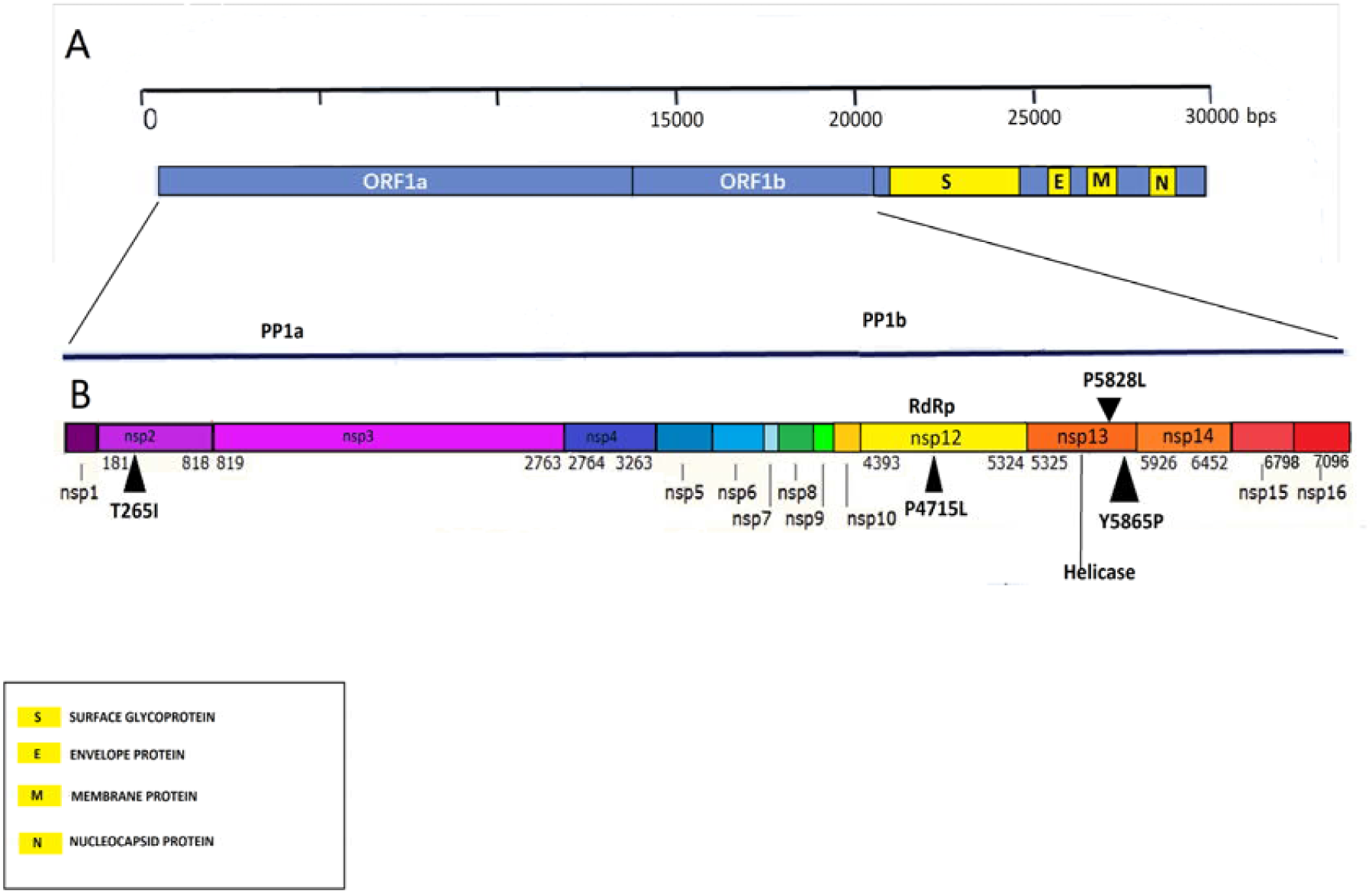
Mapping of orf1ab polyprotein and four signature mutations of USA.

Structure prediction for wild type nsps are already available in I-Tasser; which gave us the opportunity to evaluate and superimpose three altered nsp structures (nsp 2, nsp 12 and nsp 13; against said signature mutations), using UCSF Chimera (16) and PyMOL (17) for clear visualisation of the alteration (**Figure 3A**). RdRp (nsp12) is responsible for viral replication and transcription. Although SARS-CoV-2 and SARS-CoV shares only 79% genome similarity (18, 19), with regard to nsp12 sequence similarity, is comparatively much higher (98%) inferring the evolutionary significance of these conserved region (8, 20). RdRp is a major antiviral drug target (21, 22) and thus the structural alteration due to the mutant AA shall have important implications. Comparing the superimposed structure of RdRp, we have observed four small domains with the altered structure in the mutant sequence (**Figure 3B**) and these needs to be investigated further. Helicase (nsp 13) is a multi-functional protein having a zinc-binding domain in N-terminus exhibiting RNA and DNA duplex-unwinding activities with 5’ to 3’ polarity (11, 12). **Figure 3C** exhibits some important structural alteration due to those two mutations present in close proximity near the N-terminus; including a major structural alteration, where a loop or chain-like structure is transformed into a helix, found near the N-terminus of nsp 13. The length of nsp 13 or Helicase protein is 601 (5325-5925 of orf1ab polyprotein) and the above-mentioned alteration is found in the location 592-598, indicating possible variation in functional outcome.

**Figure 3.**
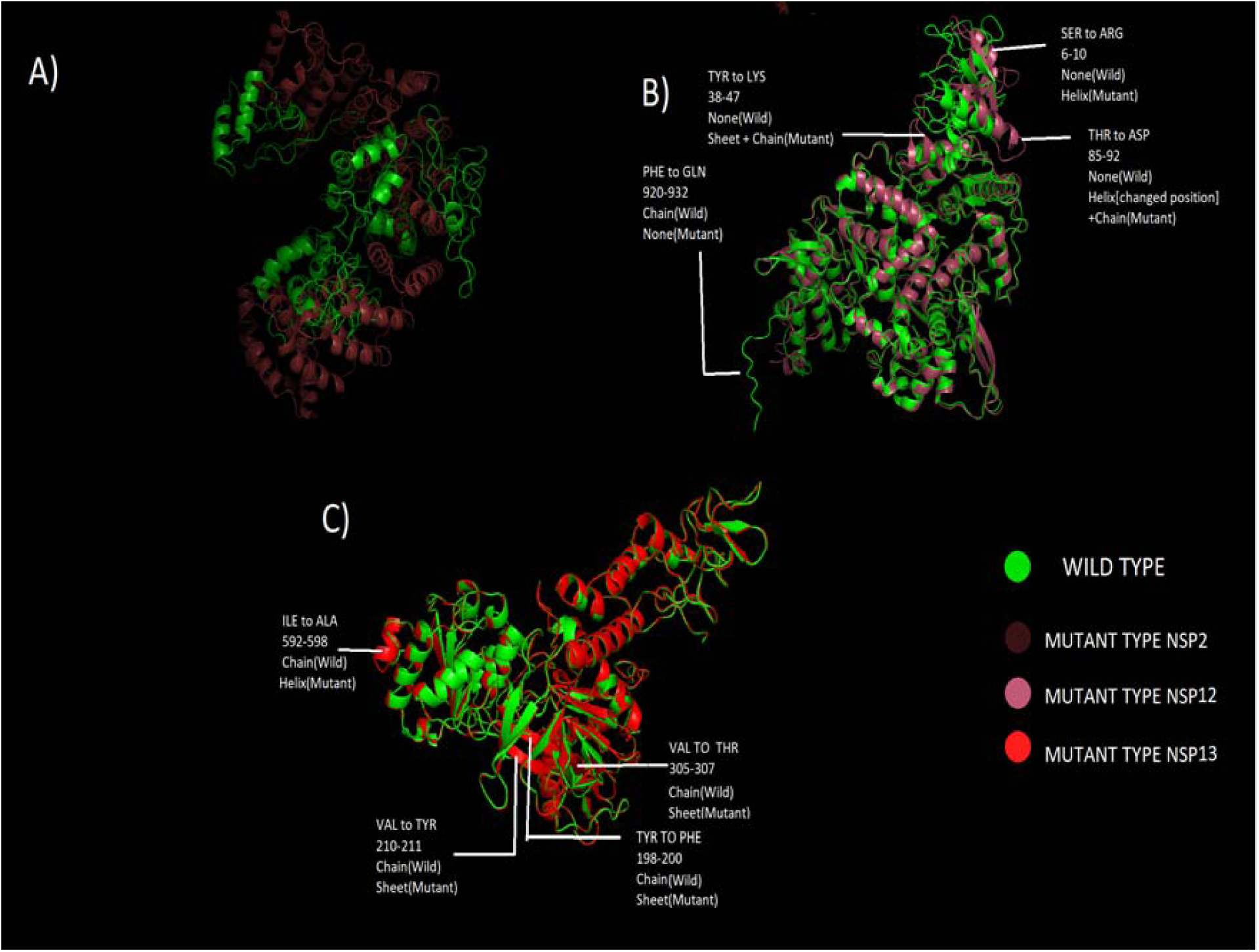
Structural comparison of nsp 2, nsp 12 and nsp 13 by superimposing the mutant structure over the wild type

## Conclusions

Orf1ab polyprotein of SARS-CoV-2 encompasses mutational spectra. Comparing to the European and Asian strain(s) we observed four characteristic mutations 265T>I (nsp 2), 4715P>L (nsp 12), 5828P>L (nsp 13) and 5865Y>C (nsp 13) which can be considered as a signature pattern for USA. It is noteworthy to mention here that 5828P>L and 5865Y>C, exclusively found in USA till now; whereas 265T>I is found in very low abundance in Asia (4.42%) and not found at all in Europe. 4715P>L is commonly found in both USA (58.1%) and Europe (51.6%) but present in low abundance in Asia(11.2%). 971P>L and 6158F>L are frequently observed in Georgia and South Carolina only representing the South-East region of the country. All of the four signature mutations caused structural alteration in their respective non-structural proteins (nsp 2, nsp 12 and nsp 13). Thereby it is essential to consider the mutational spectra while designing new antiviral therapeutics targeting viral orf1ab.

## Acknowledgement

The authors acknowledge UGC-DAE-CSR Kolkata Centre for providing fellowship to Shuvam Banerjee.

## Author Contribution

SB had the idea; SB, SS, RD, KKM did the experiments; SB analyzed the data; SB, PB interpreted the data; SS, RD, KKM, PB searched literature; SS, RD did the referencing; SB, PB wrote the manuscript; SS, RD prepared figures; PB supervised overall study.

